# Object-zoomed training of convolutional neural networks inspired by toddler development improves shape bias

**DOI:** 10.1101/2024.05.30.595526

**Authors:** Niklas Müller, Cees G. M. Snoek, Iris I. A. Groen, H. Steven Scholte

## Abstract

Convolutional Neural Networks (CNNs) surpass human-level performance on visual object recognition and detection, but their behavior still differs from human behavior in important ways. One prominent example is that CNNs trained on ImageNet exhibit an image texture bias, while humans exhibit a strong bias toward object shape. Although CNN shape bias can be increased in various ways, e.g., using data augmentation or additional training techniques, it remains unclear what causes the strong discrepancy between human and CNN object recognition strategies. Developmental research suggests that one factor driving human shape bias is that during early childhood, toddlers tend to fill their field-of-view with close-up objects. Here, we operationalize this close-up as a zoom-in on objects during CNN training which we show increases shape bias without any additional training or data augmentation. We provide further evidence for the advantage of closeup object vision by systematically manipulating the background-object ratio during CNN training, and demonstrate a strong (inverse) correlation with shape bias. Moreover, zooming-in on objects, thereby more closely emulating child vision, not only increases shape bias but also concurrently aligns classification accuracy and shape bias between humans and CNNs. Finally, we achieve a near human-like shape bias when using a developmentally-inspired background-object ratio for training and shape bias assessment. In sum, from a simple adjustment to common image datasets - zooming-in on objects - human-like shape bias can emerge. These results suggest that taking inspiration from human learning strategies is a promising avenue for building human-aligned, efficient, and more robust vision CNNs.

## 1. Introduction

Convolutional Neural Networks (CNNs) have been successful in learning and performing vision tasks such as object recognition and detection, achieving human-level or superior accuracy (Krizhevsky et al., 2012; He et al., 2016). While humans are thought to process and code information efficiently (Thorpe et al., 1996; Simoncelli and Olshausen, 2001), CNNs have been found to use shortcut learning for task completion (Geirhos et al., 2020; Hermann et al., 2023). In shortcut learning, networks exploit statistical regularities in the training data that, while predictive of the target label, may not capture the underlying conceptual relationships humans use. For instance, a network might classify images based on background elements that consistently co-occur with certain objects, rather than learning to recognize the objects themselves (Tian et al., 2021). Another prominent example, highlighting the divergence between human and CNN visual processing, is the phenomenon of texture bias (Geirhos et al., 2018). Studies have demonstrated that CNNs trained on ImageNet (Russakovsky et al., 2015) rely heavily on texture patterns when classifying objects, often failing to generalize when object shapes are preserved but textures are modified (Geirhos et al., 2018, 2020; Hermann et al., 2020; Tuli et al., 2021; Ding et al., 2022). This stands in stark contrast to human visual processing, where shape serves as a primary organizing principle for object categorization from early childhood onward (Landau et al., 1988; Cutzu and Tarr, 1997; Blanz et al., 1999; Blais et al., 2009; Ritter et al., 2017; Huber et al., 2023).

Insights from the development of human visual processing during early childhood offer promising directions for addressing these limitations. Recent research suggests that taking inspiration from the physiological development of human vision, in particular the fact that infants have low visual acuity, can help build computational models that are more aligned with humans (Jang and Tong, 2024). However, there are many other aspects of human development that have been less considered as a resource of inspiration for artificial vision models, for example, how children learn to assign object names to visually perceived inputs (Clerkin et al., 2017). By incorporating insights from developmental psychology, we may be able to guide CNNs toward learning strategies that better mirror human visual cognition, potentially leading to more generalizable and interpretable networks.

Here, our specific focus is on a set of findings in developmental psychology that show that the view of toddlers, who are learning to name objects, is frequently dominated by close-up objects (Smith et al., 2007, 2011; Bambach et al., 2018; Borjon et al., 2018). Using egocentric video recordings from both toddlers and adults interacting with objects and each other, Smith et al. (2007) identified differences in object perception between toddlers and adults, specifically demonstrating that objects frequently occupy a larger area of the visual field of toddlers, compared to adults. In contrast, large-scale image datasets commonly used for object recognition training such as ImageNet (Russakovsky et al., 2015) are often composed of images that contain a substantial amount of non-object information, i.e. background areas. In this paper, we use insights from developmental psychology to distill a training paradigm for CNNs that aims at providing more human development-like images during object recognition training, whereby training images show close-up objects. Specifically, we use bounding box annotations of various sizes to systematically reduce the background area, which thereby effectively functions as a zoom-in or close-up onto the object for which the corresponding label needs to be learned. We thereby aim to provide CNNs with the same object-dominated view as that of toddlers. By aligning human and CNN object focus in this way, we anticipate to reduce texture bias and increase object shape representation in CNNs.

Our contributions are the following. To assess the impact of object-zoomed training on shape bias in CNNs, we evaluate CNNs trained with various levels of zoom-in on images that feature conflicting cues for shape and texture (cue-conflict images), to obtain a metric for shape bias of networks (Geirhos et al., 2018). Our key finding is that CNNs that were trained on images with considerably less background information show a stronger shape bias. Thus, we show for the first time that the degree of inclusion of backgrounds during object recognition training has a direct impact on the shape bias of CNNs.

In a series of follow-up analyses, we conduct more in-depth investigations of the influence of background on CNN shape bias. First, we show that CNN shape bias cannot be studied independent from the background-object ratio of both training and cue-conflict testing images. Networks that appear to be texture-biased primarily use background information for classification (Tartaglini et al., 2022); we show here that they do not necessarily have a bias towards object texture but that the texture bias score is inflated because of the network’s usage of background information. Consistent with this idea, we show that networks that are tuned more to object information, and thus less to background information, have a higher shape bias. Second, we demonstrate that a previously reported negative correlation between CNN shape bias and test accuracy across different networks (Hermann et al., 2020) can be inverted when shape bias is assessed on cue-conflict images that are created from images containing minimal background information. The resulting positive correlation between CNN shape bias and test accuracy then closely resembles the pattern observed when evaluating shape bias and accuracy of humans.

Finally, we find that a near human-like shape bias is achieved when using a background-object ratio that closely resembles the object-dominated view of toddlers. Excessive zoom-in which leads to object cropping that disrupts the global object shape again decreases the shape bias of networks. This suggests that there is a specific optimal range with which CNNs can be trained to achieve high shape bias. Specifically, using a zoom-in that removes the maximum amount of background information while keeping the global object shape intact leads to simultaneous alignment of shape bias and test accuracy between humans and CNNs.

Overall, we present empirical findings of human-like shape bias in CNNs when reducing the background-object ratio during training, inspired by object zoom-in behavior observed during human development. Due to this reduction of background information, training images primarily contain object-related information, which improves the object classification supervision signal, leading to a developmentally inspired, biologically plausible alignment of human and CNN object classification strategies.

## 2. Related Work

The strong texture bias of CNNs was first demonstrated by Geirhos et al. (2018) using images that exhibit conflicting cues for shape and texture. Images were designed to depict the shape of an object superimposed with the texture of another, e.g. the shape of a cat with the texture of an elephant skin. Networks primarily classify these images with the corresponding texture label, while humans in most cases indicate the shape class to be most descriptive of the object in the image. This striking difference between shape-biased humans (Landau et al., 1988) and texture-biased CNNs has not only sparked a debate on whether CNNs are suitable models of human cognition (Bowers et al., 2023) but also led to a variety of approaches aiming to increase CNN shape bias, which we outline in more detail below. Finding ways to increase shape bias is worthwhile for both computer scientists and cognitive neuroscientists as it could lead to more robust and generalizable network representations and to a better understanding of the origins of these biases in both humans and networks. However, it is still unclear what aspects of the training of CNNs matter for developing object feature biases. Prior work suggests that e.g., learning strategies play a crucial role in the development of categorization biases (Hermann et al., 2023).

Various methods have been proposed that successfully increase CNN shape bias. This includes modifying the architecture by using convolutional networks with larger convolutional kernels (Ding et al., 2022), Vision Transformers (Tuli et al., 2021), or generative classifiers (Jaini et al., 2023). Although both Transformer-based models (Dosovitskiy, 2020) and generative classifiers (Jaini et al., 2023) have been shown to exhibit high task performance and increased shape bias, they seem to be solving the task of object recognition using an alternative processing strategy than the brain or convolutional architectures (Tuli et al., 2021). An alternative strategy, in which the model architecture remains fixed, but the training images are augmented, has also been shown to shift network bias toward shape (Geirhos et al., 2018). By making texture features less reliable across repetitions of the same image as well as across instances of the same object class, augmentations can be used to directly reduce the bias towards texture (Hermann et al., 2020). However, it remains unclear whether networks merely become worse at classifying texture (thus resulting in an increase of shape bias) instead of improving classification accuracy on object shape. Augmenting images to express object shape more vigorously, by exploiting segmentation masks or edge information, has also been found to increase shape bias (Lee et al., 2022; Gowda et al., 2022).

Although these approaches have achieved success, they either manipulate the model architecture, implying that texture bias is inherent to certain CNN architectures (e.g. the commonly used ResNet50, He et al., 2016), or manipulate the input data by using augmentations that are explicitly aimed at enhancing the learning of shape features. Humans certainly encounter objects in diverse settings and from changing viewpoints and scales, and the human brain appears to have dedicated mechanisms for computing shape features (Ayzenberg and Behrmann, 2022). However, many of the augmentations that are typically applied during network training are not plausible from the perspective of human learning (e.g. random grayscale or color jitter). Thus, both architectural changes and strong input manipulations fail to draw clear parallels to the origins of human shape bias. Here, we distill a training paradigm for CNNs directly from the current understanding of how shape bias develops in humans.

Recent work shows that taking inspiration directly from human visual development in training CNNs can increase shape bias by applying blur during training (Jang and Tong, 2024). The low visual acuity of infants is thought to be well resembled by blurred images in which low spatial frequencies are the dominating visual features which are sufficient for the identification of object shapes. However, toddlers learn to verbalize and to associate objects with names at later stages of human development, when high acuity vision has fully developed. During this period single objects frequently dominate the view of toddlers and can be seen as a closer zoom-in of their physical environment (Smith et al., 2007, 2011). This stage of human development can be thought of as comparable to training CNNs to assign category labels to objects. A direct comparison of training CNNs on training images that resemble either the view on objects of adults or of toddlers has shown that the CNNs trained on the toddler training set showed increased generalization performance (Bambach et al., 2018; Aubret et al., 2022). However, whether mimicking this toddler-like object learning can account for the differences in shape bias between humans and CNNs has not yet been examined. We thus asked whether taking inspiration from this stage of human development by training CNNs on close-up objects will increase their shape bias and thus lead to improved human alignment.

## 3. Methods

### 3.1 Image datasets

To examine how the background-object ratio of training images affects the shape bias of CNNs, we use two commonly used image datasets, MS-COCO (Lin et al., 2014) and ADE20K (Zhou et al., 2017), as well as our own Open Amsterdam Data Set (OADS) (see e.g. Mueller et al., 2023, 2024a,b). Each dataset features different annotations and thus offers different ways of controlling the background-object ratio, which are explained below. In this paper, we use the term shape bias, instead of texture bias or texture-shape bias, to increase the ease of comparisons to shape-biased humans. We note that the reported texture bias in other literature can directly be derived by calculating 1− ”*shape bias*”.

First, we used publicly available object bounding box annotations for MS-COCO images to create object crop datasets. We varied the amount of image area surrounding the object bounding box that was included in the object crop, from 0% (tight) to 80% (narrow) and 150% (wide) of additional surrounding image area that was included (see Fig. 1a).

**Figure 1:**
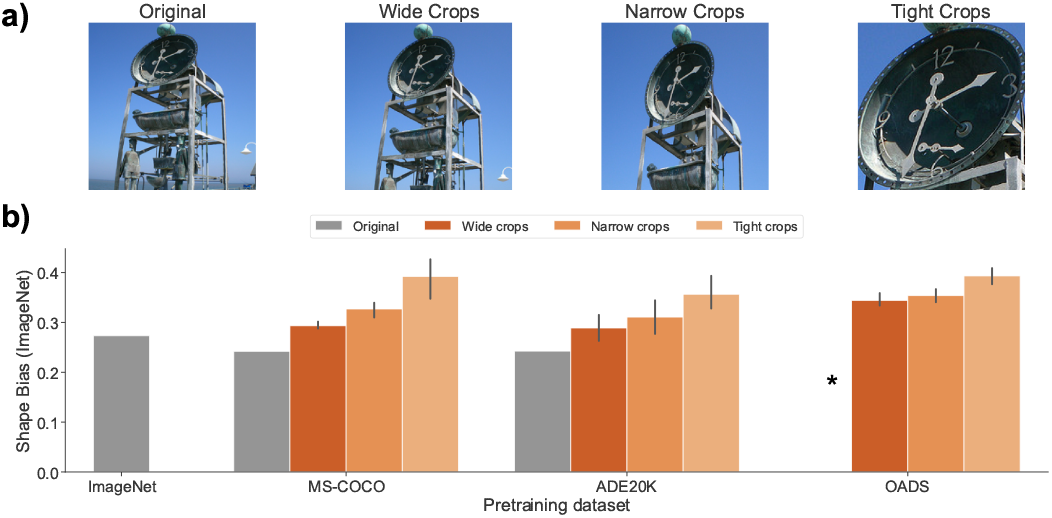
Zoomed-in object training increases ImageNet shape bias. **a)** Examples of original images and wide, narrow, and tight crops. Example image is from the MS-COCO dataset. **b)** Shape bias on ImageNet cue-conflict images of networks trained on three different datasets: MS-COCO (Lin et al., 2014), ADE20K (Zhou et al., 2017), and the Open Amsterdam Data Set (see Methods section 3.1), a custom dataset consisting of ultra high-resolution scene images, featuring manually annotated object bounding boxes and object category labels. Using OADS allows for close control of object crops both during training and for creating a new cue-conflict test set, independent from the original ImageNet benchmark. We trained networks on either full images (gray), or on object crops created using bounding boxes (orange shades, see a)). Per dataset, we show the average shape bias across four ResNet50 instances using different random seeds. Error bars indicate the 95% confidence interval. (* there is no model equivalent to trained on original images for OADS). This shows that shape bias increases when reducing the amount of background included in training crops in all datasets.

Second, we created object crop datasets for ADE20K images using publicly available object segmentation annotations, from which we created object bounding boxes with the minimal possible rectangular area that subtends the fully segmented object. Following the above strategy of creating tight object crops, we then created three ADE20K object crop datasets with varying degrees of bounding box tightness.

Last, we used our custom OADS dataset consisting of 6130 uncompressed scene images with a resolution of 5496×3672 pixels. For each scene image in OADS multiple objects were manually annotated with a bounding box and one of 16 object class labels (see table A.2) resulting in 64.683 annotated objects. We used these bounding boxes to create object crop datasets, again using three degrees of crop tightness. OADS was additionally used to create a new cue-conflict test set (see section 3.3) to assess texture-shape bias (see section 3.5) independent from classification ability on ImageNet.

### 3.2 Training convolutional neural networks

For the nine object crop datasets (wide, narrow, and tight crops for MS-COCO, ADE20K, and OADS, respectively) described above, we trained separate ResNet50 (He et al., 2016) instances on object classification for 30 epochs. We repeated model training for each dataset 4 times using different random seeds. Number of images in training and validation datasets, number of classes, as well as validation accuracy are provided for each dataset in Table A.3. We focus on models that inherently process local spatial information hierarchically (as opposed to, e.g., Transformer architectures) as this is a core principle of the human visual system (Hubel and Wiesel, 1962; Dumoulin and Wandell, 2008).

To test if our results are task- and dataset-specific, we also tested networks pretrained on an object detection task on standard MS-COCO images (Long et al., 2015), on a scene-segmentation task on standard ADE20K images (Cheng et al., 2021), and networks trained on object classification on standard ImageNet images. For the object detection and scene segmentation networks, we only used their ResNet50 backbone and replaced their task-head with a 2-layer fully-connected object-classification head. We note that due to the large size of the OADS images, there is no model trained on full scene images (indicated by the asterisk in Fig. 1a).

Since the shape bias benchmark using cue-conflict images developed by Geirhos et al. (2018) (see section 3.3) is based on images and image classes from ImageNet, evaluating shape bias in our networks on this benchmark required finetuning on ImageNet. We finetuned each pretrained model on a subset of ImageNet for 15 epochs (see Table A.4 for test accuracies after finetuning). The subset consisted of all images of those 223 classes that were used to create the ImageNet cue-conflict images in Geirhos et al. (2018) (see Table A.3 for number of images in training and test set; training set was used for finetuning). Additionally, to test whether finetuning itself has an effect on shape bias, we varied the amount of parameters that were frozen during finetuning and either only trained the classification head (<1% parameters finetuned), the last residual block (∼33%), or the full model (100%).

All custom networks were trained using an Adam optimizer (learning rate: 0.001) on two NVIDIA A100 and 18 cores, with randomly chosen train and test splits (reported in table A.1). A typical training run for object classification on ADE20K was 13 hours, for MS-COCO 3 hours, and for OADS 3 hours. A typical finetuning run on ImageNet took 2 hours.

### 3.3. Shape bias assessment

Shape bias assessment for ImageNet was performed on all 1280 cue-conflict stimuli taken from the Geirhos et al. (2018) benchmark. Briefly, images in this benchmark consist of a subset of ImageNet images that are manipulated to contain conflicting cues for object shape and object texture combined using neural style transfer (Gatys et al., 2016). The intention for these images is that they express the shape of one object and the texture of another (e.g. a shape of a cat with the texture of an elephant skin). Evaluating the classification performance of networks on these images therefore reveals a preference for either texture or shape (e.g. if the model classifies the image as a cat, it is shape-biased, whereas if it classifies it as an elephant, it is texture-biased).

For each cue-conflict image, CNN decisions are based on the class associated with the maximally activated output node. Following Geirhos et al. (2018), shape bias was calculated as the number of shape decision over the sum of shape and texture decisions. Additionally, cue-conflict accuracy was calculated as the sum of correct decisions for the shape and the texture class over the total number of images. For our last experiment, we computed accuracy on cue-conflict images separately, for counting either only the shape class as correct or only the texture class as correct, and then dividing the sum of correct decisions by the total number of images.

To test shape bias independently from ImageNet classification ability, we additionally created a new set of cue-conflict images for OADS using the same methodology and hyperparameters as for ImageNet cue-conflict stimuli from (Geirhos et al., 2018). From a set of tightly cropped OADS object images for which the standard OADS-trained ResNet50 had 100% classification accuracy, we randomly drew 62 images and performed neural style transfer for each pair of images. We randomly selected 880 from the resulting 3844 images, while maintaining the balance between number of images per class. We created 9 aggregate classes (see italic classes in table A.2) merging similar subclasses, similar to (Geirhos et al., 2018). OADS cue-conflict images were created with a resolution of 400×400 pixels.

### 3.4. Cue-conflict image masking

To test if our method of training on tight object crops reduces network tuning to background features, we used the approach from (Tartaglini et al., 2022): we assessed shape bias and accuracy of CNNs separately for cue-conflict images where either the background was masked or the object was masked (see Fig. 2a), leaving the respective other area fully intact. Masking was done using silhouette images from Geirhos et al. (2018) and entails setting all pixels included in the mask to zero while leaving others unchanged.

**Figure 2:**
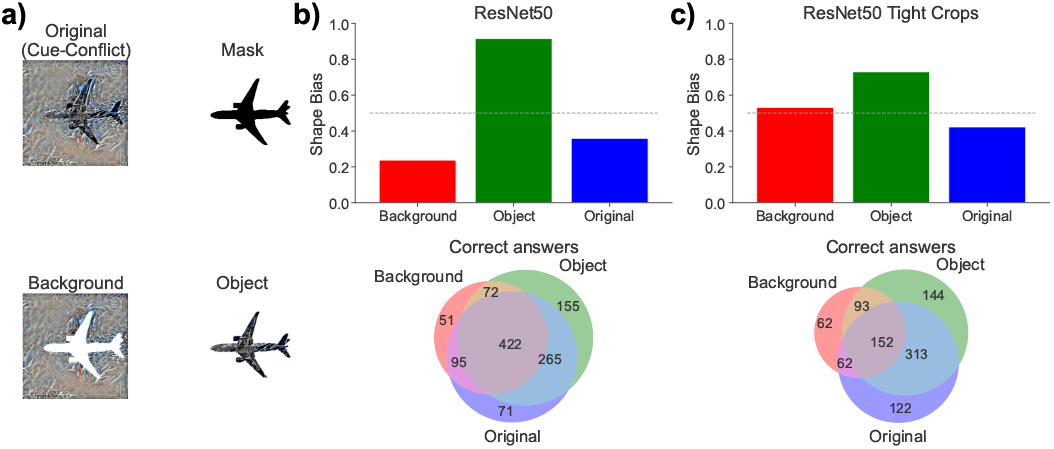
Texture bias on ImageNet results from networks using background information. **a)** Example of an original cue-conflict image, the associated object silhouette (“mask”) that was used to mask the object, the original cue-conflict image without the masked object (“background”), and inversely, the original cue-conflict image while masking out the background. **b)** Shape bias for the default ImageNet-trained ResNet50 when using the original cue-conflict images or the object or background masked images (top). Venn diagram showing the number and overlap of correct answers for each masking condition (bottom). c) Same as b) but for a model pretrained on MS-COCO tight object crops and fully finetuned on ImageNet.

### 3.5. Human behavioral experiment

We performed a psychophysical experiment on human participants in the lab using the OADS cue-conflict images to assess human shape bias on the newly created test set. Collection and use of human behavioral data was approved by the Ethics Review Board of the University of Amsterdam. We replicated the experimental paradigm from Geirhos et al. (2017) as much as possible. Specifically, 14 participants (Mean age: 19.8 + −2.5, 10 female) each performed 880 trials, where each trial consisted of the presentation of a central fixation cross for 300 ms, followed by the presentation of a randomly drawn, non-repeating OADS cue-conflict image for 200 ms, followed by a pink noise mask for 200 ms, followed by a 2500 ms time window to give a response in the form of a mouse click on one of 9 object class names presented on the screen. The 27” screen had a resolution of 2560 × 1440 pixels and images were presented in the center of the screen with a resolution of 400 × 400 pixels and thus subtended an area of 8.5 × 9.0 degrees of visual angle with a viewing distance of 63 cm. Participants were instructed to choose the object class name that best describes the previously seen object by clicking on the class name on the screen. Participants were introduced to the set of classes before the start of the experiment, and order and arrangement of class names on the screen did not change throughout the experiment. Further, performance of participants (calculated as accuracy, counting the selection of both the texture and the shape class as correct) was presented to the participant every 5 trials. After every 50 trials, participants were given a break for at least 30 seconds and self-initiated the next set of trials. Participants were rewarded with 12.5 euros or an equivalent of 1.5 research credits. We excluded one participant who had an accuracy of more than two standard deviations below the population mean.

## 4. Results

To investigate the relation between background-object ratio and shape bias of CNNs trained on object recognition, we manipulated object crop tightness during both training and testing, thereby systematically varying the relative amount of object and background information accessible to the networks. First, we show that ImageNet shape bias robustly increases when using tighter object crops during pre-training on a variety of datasets. Second, by separately evaluating biases for object and background, we highlight that high texture bias on ImageNet can result from networks using background information instead of using object texture. Next, we show that a previously reported negative relation between shape bias and task accuracy (Hermann et al., 2020) also applies when incrementally finetuning pre-trained networks on ImageNet. We further leveraged our custom high resolution scene dataset to demonstrate that this negative relation can be inverted when images used to evaluate shape bias contain minimal background information. Finally, we show that it is possible to achieve a near human-like shape bias by simply using an optimal background-object ratio that reflects the close-up view of child vision.

### 4.1 Training on tight object crops increases CNN shape bias on ImageNet

First, we varied the amount of background information in training images by manipulating the size of object bounding boxes on a variety of datasets. Specifically, we used the ImageNet-based benchmark proposed by Geirhos et al. (2018) for testing shape bias using cue-conflict images (see Methods section 3.3) to assess shape bias of CNNs pre-trained on object datasets derived from MS-COCO, ADE20K, and a custom, high-resolution scene dataset OADS (see Methods section 3.1). For each dataset, multiple versions were created, varying the degree of crop tightness, ranging from maximally tight object crops (0% surroundings), over narrow crops (80%) to wide crops (150%) (see Fig. 1a). We additionally included a comparison with a baseline model trained on original ImageNet images, as well as publicly available networks pre-trained on full scenes (object detection task for MS-COCO and object-scene segmentation for ADE20K) as they contain the maximum amount of background information possible.

Results show that shape bias increases systematically with crop tightness for all three datasets (see Fig. 1b). Moreover, networks trained on full scenes (gray bars in Fig. 1, referred to as ”Original”) have the lowest shape bias, further highlighting the detrimental effect of background information on shape bias. Importantly, these original networks, which were pretrained using a segmentation or detection task objective, also show no increase relative to original ImageNet-trained model (see gray bars in Fig. 1b), indicating that the observed increase with crop tightness is not due to a change in task objective. Together, these results suggest that there is a direct effect of including background information during training on the shape bias of CNNs.

### 4.2 CNNs trained on tight object crops rely less on background features

To better understand why reducing background results in increased shape bias, we inspected the classification strategy of the networks in a setting in which object and background are explicitly disambiguated. Previous work (Tartaglini et al., 2022) has shown, by applying object masking (see Fig. 2a; for more details see section 3.4), that using cue-conflict images alone cannot differentiate between a bias towards object texture or background. Specifically, when masking out the background, this leaves the object area of the original shape image superimposed with the texture of the texture image. In this condition, a truly texture-biased model should still classify the image according to the texture category. However, they found that under these circumstances the model was in fact strongly shape-biased, suggesting that the model’s texture bias actually arose from the usage of background information (see Fig. 2b).

If a low shape bias indeed reflects a bias toward background information, then our method of removing background information from training images should reveal an increased usage of object features (texture and shape) in this setting. Such an increased usage of object features should translate into a low classification accuracy when the object is masked out, showing that the model does not base its predictions on the image background. Indeed, we find that a model trained on tight object crops of MS-COCO gives fewer correct answers on cue-conflict images when only presented with the object background (see red circle in Venn diagram in Fig. 2c). This shows that this model does not rely as strongly on background information.

Next, to test whether this model, that is trained on tight object crops, instead relies more on object features, we presented it with cue-conflict images containing only the object while the background is masked out. Now, the model predicts correct classes well, showing that it uses object features for classification, and has a high shape bias (see green bar in Fig. 2c), showing that it uses object shape over object texture.

These results directly confirm that our method of object zoom-in reduces reliance on background features, which translates into a reduced texture bias on ImageNet. It remains unclear, however, whether the usage of background information that leads to a high texture bias, stems from the assessment of shape bias on ImageNet cue-conflict images or from the training on ImageNet images. In the following section, we therefore examine the impact that finetuning on ImageNet has on the shape bias of CNNs that are pre-trained on tight object crops.

### 4.3. Training on ImageNet induces low shape bias

The results above reveal a direct link between the amount of background included in training images and shape bias on the conflict benchmark by Geirhos et al. (2018). However, the gains in shape bias due to background reduction (Fig. 1B), were relatively modest, not yet approaching human-level shape bias (which tends to average around 0.8 or 0.9, Geirhos et al. (2018)). Cue-conflict images from this benchmark depict objects that have the shape and texture of ImageNet classes, and therefore networks need to be able to classify images according to the ImageNet classes in order to establish shape bias. Thus, shape bias assessment using this benchmark necessitates some amount of finetuning on ImageNet. Here, we asked whether this finetuning procedure potentially diminished the effects of object crop training, thereby contributing to the relatively modest increase in shape bias in our CNNs. To test this, we systematically evaluated the impact of different amounts of finetuning on network shape bias. For networks pre-trained on object classification on MS-COCO, ADE20K, and OADS, instead of finetuning the full model on ImageNet as in the previous section, we now kept an increasing number of parameters (see Methods section 3.2) fixed during finetuning, thereby explicitly maintaining the features from the pre-training.

Fig. 3a) shows that shape bias gradually and systematically decreases with increasing amount of finetuning (see colored clusters). Finetuning only a small number of parameters (<1%, blue points) yielded networks that are shape-biased (but have low test accuracy, due to the low amount of finetuning). Increasing the number of parameters that were finetuned (∼33%, orange points) until all were included (100%, green points) led to a decrease in shape bias coupled with a simultaneous increase in test accuracy. Full finetuning on ImageNet led to comparable shape bias and test accuracy as training networks on ImageNet from scratch (see red point). Hence, even within the same model there is a strong negative correlation between test accuracy on ImageNet and shape bias (*r* = −0.80) driven by the amount of parameters that were finetuned on ImageNet. This correlation is even more pronounced when evaluating cue-conflict accuracy (*r* = −0.84, see Fig. 3b), i.e. the sum of correct shape and texture decision over the total amount of cue-conflict images (see section 3.3).

**Figure 3:**
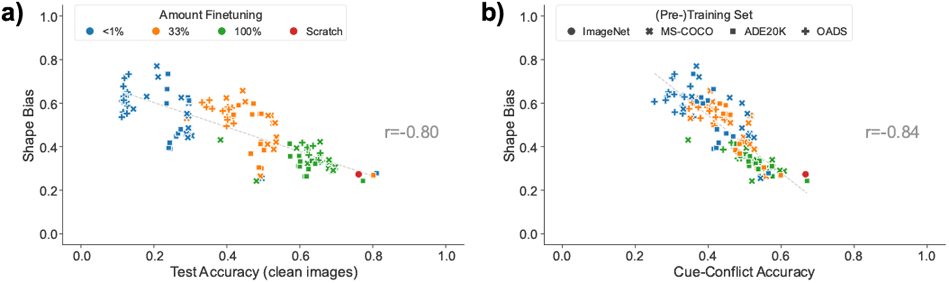
Shape bias decreases with increasing accuracy on clean and cue-conflict images. **a)** Test accuracy (default ImageNet test set) vs. shape bias and **b)** cue-conflict accuracy vs. shape bias for networks pretrained on various datasets (markers) and with varying amounts of parameters (colors) finetuned on ImageNet. Shape bias decreases with test accuracy on default test images and with accuracy on cue-conflict images (used for measuring shape bias). A network with high accuracy and high shape bias would then be human-aligned.

These results show that our previously demonstrated benefits of pre-training on tight object crops (see Fig. 1) systematically diminish with increasing amounts of finetuning on ImageNet. Our pretrained networks exhibit a high, near human-like shape bias when only minimal finetuning on ImageNet is performed, but this high shape bias trades off against a low performance accuracy on classification. This suggests that, for the models we tested here, simultaneous high shape bias and high accuracy cannot be achieved on ImageNet. However, higher gains of shape bias can potentially be achieved when testing shape bias on cue-conflict images created from other datasets. We explore this direction in the next section.

### 4.4. Training on tight object crops resolves the accuracy vs. shape bias tradeoff

So far, we have found that the classification strategy of CNNs is influenced by the amount of background information in training images, as a gradual removal of background information led to a gradual increase in shape bias (section 4.1). However, this increase in shape bias comes with a substantial drop in model accuracy (section 4.3), presumably due to the finetuning on ImageNet leading to a strong sensitivity to background features (section 4.2). In the following, we show that this tradeoff is resolved when background information was reduced not only in the training images but also in the cue-conflict images used to evaluate shape bias. Specifically, we used our custom, high resolution image dataset OADS, featuring manually annotated object bounding boxes, to create a new cue-conflict dataset based on tight object crops: OADS cue-conflict (see Methods section 3.1 for details). These OADS cue-conflict images contain minimal background information and thus allow disentangling the model’s usage of object texture and background information.

First, to assess shape bias of humans on the new OADS cue-conflict dataset, we ran a psychophysical experiment (see section 3.5). We found that shape bias was higher than 0.8 for all participants, comparable to the human shape bias on ImageNet cue-conflict images (see left panel in Fig. 4a). Moreover, as in the previous benchmark, human participants showed a positive correlation between cue-conflict accuracy and shape bias in the OADS cue-conflict task.

**Figure 4:**
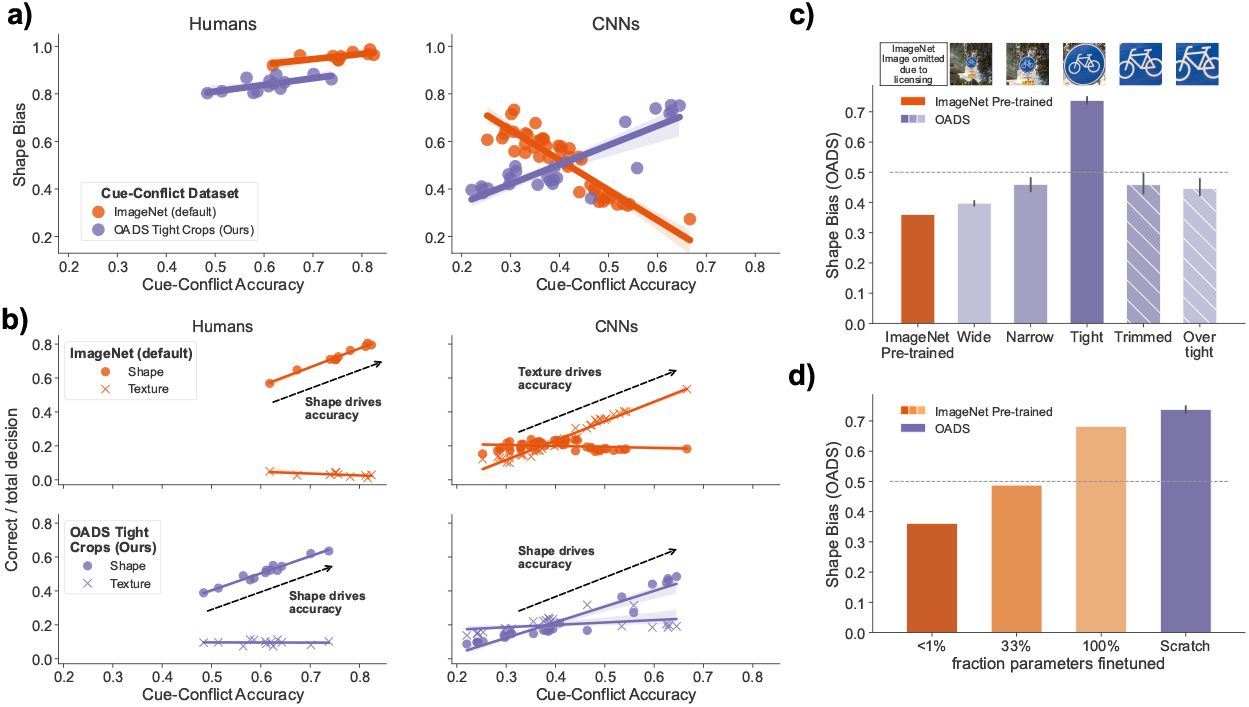
Shape bias assessment is human-aligned when using tightly cropped cue-conflict images. **a)** Accuracy (on cue-conflict images) vs. shape bias of individual humans and individual CNNs for ImageNet (orange) and OADS (ours; purple). OADS cue-conflict images are created from tight object crops. Circles show accuracy and shape bias for individual humans and CNNs and solid lines show the best fitting regression line. **b)** Correct decisions over the total number of decisions calculated for considering the shape class the correct class (circles) or the texture class the correct class (crosses), for humans (first columns) and CNNs (second column) assessed on ImageNet cue-conflict images (top row) and OADS cue-conflict images (bottom row). Individual points and solid lines as in a). **c)** Shape bias measured on OADS cue-conflict images for a model pre-trained on ImageNet (orange) and for networks trained on varying crop tightness on OADS (purple), including crops that are smaller than the object bounding box and thus remove parts of the global object shape. **d)** Shape bias for networks pre-trained on ImageNet and finetuned on OADS while varying the number of parameters that are finetuned (orange), and a model trained from scratch on OADS tight crops (purple).

Next, we use the same set of CNNs pre-trained on object recognition as before (Figs. 1 and 3), but now explicitly compare their shape bias after finetuning on ImageNet (see orange dots in right panel in Fig. 4a) with their shape bias obtained after finetuning on OADS (if they were not pre-trained on OADS; see purple dots). Results show that finetuning on OADS gives rise to the opposite correlation between shape bias and accuracy. As previously highlighted, ImageNet shape bias decreases with increasing cue-conflict accuracy, while the same networks finetuned on OADS show an increasing shape bias with increased cue-conflict accuracy.

To better understand these diverging trends between accuracy and shape bias, we evaluated the shape accuracy and texture accuracy separately, i.e. counting the number of correct shape decisions (see circles in Fig. 4b) or the number of correct texture decisions (see crosses in Fig. 4b) over the total number of images, respectively. For humans (first column) texture accuracy stays constant across participants with varying cue-conflict accuracy while shape accuracy increases with increasing cue-conflict accuracy, for both cue-conflict benchmarks. For CNNs, there is a categorical difference between cue-conflict benchmarks: on ImageNet, networks’ accuracy increases because more images are correctly classified by their texture. Contrarily, on OADS, networks’ accuracy increases because more images are correctly classified by their shape. The positive correlation between accuracy and shape bias with OADS finetuning now accurately resembles the pattern observed in human participants: like for humans, accuracy of networks that have a high shape bias is driven by their classification accuracy of shape, while the accuracy of networks that have a low shape bias is driven by their classification accuracy of textures.

We furthermore find that the effect of training with object zoom-in on shape bias is even more pronounced when both training and testing networks on OADS objects. An OADS pre-trained network with tight crops exhibits a shape bias that is twice as large as that of a standard CNN pretrained on ImageNet (and then finetuned on OADS) (Fig. 4c). Interestingly, however, continuing to crop images does not continue to result in even higher shape bias. To demonstrate this, we extended the previous experiment and increased crop tightness even further, removing additional background information but crucially also removing parts of the global object shape. We expected that a model trained on these trimmed images would show a lower shape bias and instead learn to classify objects according to their texture. Indeed, we find that networks trained on trimmed or over tight object crops again have a lower shape bias (see Fig. 4c) two right-most purple bars). Thus, this shows that continual increases of crop tightness do not lead to a monotonous increase of shape bias, and rather suggests that there is an optimal background-object ratio instead, that leads to high shape bias in CNNs and thus to an increased alignment with humans.

Finally, to showcase that a small adjustment to training images can have a large impact on the object classification strategy of (even pre-trained) CNNs, we performed the inverse experiment as in Fig. 3: we took an ImageNet pre-trained network, which has a low shape bias and uses background features for classification, and assess its shape bias after finetuning on tightly cropped OADS objects. As before, we systematically increased the number of parameters that are finetuned on OADS. We find that shape bias increases with the amount of parameters finetuned on OADS (see Fig. 4d), approaching, yet not surpassing, the high shape bias of networks trained on OADS from scratch. This shows that small amounts of finetuning on data that facilitates learning to classify objects using object features instead of background features is a simple solution for increasing the shape bias of CNNs, and demonstrates that the characteristics of the training data has a large influence on the task strategies of CNNs.

In sum, these results show that, for the models we tested, behavioral alignment on shape bias between humans and CNNs can be achieved through the removal of background information in both training and cue-conflict images, but not when using the original ImageNet dataset and benchmark. This suggests that a high texture bias is not an inherent property of CNNs but can instead be mitigated when using images that have a more optimal background-object ratio, providing a more informative training signal when training CNNs to classify objects.

## 5. Discussion

The divergence between human and machine visual processing has emerged as a critical focus in both computer science and cognitive science. While CNNs have become the prime candidates for modeling human cognition and visual processing, they have been found to exploit statistical regularities in the training data that allow them to accurately perform their task while using information that humans typically do not use. This behavior is termed shortcut-learning and expresses itself for example in CNNs’ reliance on shape versus texture: for humans, objects are typically identified by their distinct shape while CNNs have been shown to use image texture for classifying objects (Geirhos et al., 2018). This ”texture-shape bias” disparity has attracted significant attention across multiple fields: in computer science and representation learning, researchers have explored architectural and training innovations to address this gap (Hermann et al., 2020; Geirhos et al., 2021; Li et al., 2024; Orhan and Lake, 2024) while neuroscientists have investigated its implications for understanding human visual processing (Kim et al., 2019; Jagadeesh and Livingstone, 2024; Subramanian et al., 2024). Here, we aim to resolve this disparity by drawing inspiration from a less often considered source: early human visual development. Given prior findings showing how toddlers naturally interact with objects during critical learning periods — specifically their tendency to view objects up close while learning to name objects (Smith et al., 2007, 2011; Clerkin et al., 2017; Bambach et al., 2018; Borjon et al., 2018) - we here explore how training neural networks with images of close-up objects influences shape bias and recognition accuracy. We demonstrate that such a small, yet ecologically motivated change to training images has a large effect on the behavior of CNNs and their alignment with humans.

### 5.1. CNN shape bias is directly linked to training data characteristics

Our research suggests that the low shape bias prevalent in CNNs may be more a consequence of training data characteristics than inherent to the network architecture. Through systematic manipulation of background-object ratios in three image datasets (MS-COCO, ADE20K, OADS), we have uncovered several new insights into how shape bias is linked to training data characteristics.

First, we showed that stepwise reductions of the amount of non-object-bound (background) information leads to the emergence of a stronger shape bias in CNNs. The reduction in background information matches how humans naturally process visual information during development. However, we also showed that stronger cropping that disrupts global object shape led to a decrease in shape bias. Such removal of object shape may also be an (unintended) side effect of data augmentation strategies commonly applied in CNN training - our results suggest that this cropping may be a driving factor in models developing a stronger bias towards texture. Interestingly, there seems to be an optimal background-object ratio when aiming to increase the shape bias of CNNs. Training CNNs using an optimal background-object ratio specifically matters for reducing shortcut-learning as CNNs have been found to rely more on available than on relevant visual features (Hermann et al., 2023). This demonstrates the importance of dataset quality when building computational models of human visual processing.

### 5.2. Disentangling the origin of texture bias

Second, we expanded on previous work that showed that the ImageNet cue-conflict images (Geirhos et al., 2018) do not allow to distinguish between an object texture bias and a background bias, with the backgrounds leading to an inflated texture bias and with that the notion that CNNs are generally biased towards object texture (Tartaglini et al., 2022). Here, by assessing not only shape bias but also cue-conflict accuracy, we showed that networks predict object classes based on information presented in the background of cue-conflict images. As a consequence, these networks show a background bias that manifests itself as a low shape bias when using the ImageNet cue-conflict benchmark. Our method of removing background information during training of CNNs alleviates this inflated texture bias as networks do not use background features for their object classification strategy but instead base their object classification decisions on object shape information. Interestingly, we found no difference in shape bias between models trained on original images on different task objectives, including object classification, object detection, and scene segmentation. Specifically, models that segment objects from their background (object detection and scene segmentation) do not develop a higher shape bias than models trained on standard ImageNet object classification. This suggests that the available information in training images has a stronger influence on the task strategy than the task objective itself, highlighting yet again, the susceptibility of CNNs to shortcut-learning.

### 5.3. Limitations of ImageNet in assessing human-machine alignment

Our demonstrated gain in shape bias when training on tight object crops systematically diminished the more networks were finetuned to ImageNet images and classes. We quantified this decreasing trend by looking at the correlation between ImageNet test accuracy and shape bias of various pre-trained models that were more or less finetuned on ImageNet. This analysis yielded three significant insights for comparing CNNs and humans: first, the correlation between ImageNet test accuracy and shape bias was negative for CNNs, while this correlation was positive for humans. Second, we showed that these diverging trends of CNNs and humans can be aligned when images that contain minimal background information are used for training of CNNs as well as for assessing shape bias for CNNs and humans. Third, we showed that the positive correlation for humans is exclusively driven by an improving performance to classify images by their shape, i.e. participants with an overall higher accuracy were only better on classifying images by shape but not by texture. This shows that humans really based their object classification ability on object shapes. Contrarily, when assessing networks on ImageNet cue-conflict images, high performing networks are better at classifying images by their texture while their number of shape decision remains constant. Distinguishing between correct shape decisions and texture decisions is important, as shape bias could also increase by a mere decrease in number of correct texture decisions, while, as we show here, for increased alignment with humans, an increase in shape bias should be driven by an increasing number of correct shape decisions. Networks assessed on OADS cue-conflict images indeed exhibit this behavior: like humans, better performing networks more often correctly classify shapes.

Last, we demonstrated that human-like shape bias is attained when CNNs are trained in a developmentally inspired manner. Specifically, maximal removal of background information while preserving the global object shape in training images led to the highest shape bias. Finetuning an ImageNet pre-trained network using those same object-focused image led to systematic increases of shape bias even when only finetuning small amounts of parameters. Thus, even a strongly texture-biased network can be turned shape-biased using object-zoomed training.

### 5.4. Limitations and future directions

The conclusions of this paper are limited by the CNN architectures and datasets used for CNN training. Specifically, we manipulated the amount of background information in image datasets that feature high quality object bounding box annotations (MS-COCO, ADE20K, OADS) and then finetuned networks on ImageNet to assess shape bias. The optimal pipeline would be a direct manipulation of ImageNet images, as well as ImageNet cue-conflict images. However, bounding box annotations for ImageNet only exist partially and are of low quality; a direct test on ImageNet would require running new behavioral annotation in order to obtain such high-quality bounding boxes, which was out of scope for the present study. We explicitly exclude Transformer architectures from this experiment as their way of processing visual information as cross-attended patches is unlikely to match the local processing hierarchy of the human visual system. Nevertheless, a future extension of this work could examine the effect of object zoom-in on Transformer architectures. The main focus of this study was to unravel how CNNs develop different object feature biases. However, this constitutes only one aspect of evaluating the behavioral alignment between humans and CNNs. Other behavioral benchmarks, e.g. error consistency (Geirhos et al., 2017), have been proposed as a more holistic comparison. Unfortunately, such an extensive benchmark for evaluating behavioral alignment between humans and CNNs only exists for ImageNet images and therefore presents similar problems as we have outlined in this work. Finally, a crucial difference between current models and humans is that models only statically perceive visual information and are not capable of interacting with their visual input, while humans gain rich information by manipulating objects (Smith, 2005) and repositioning themselves in their environment (bon Hofsten and Rosander, 1996; James et al., 2001; Smith and Gasser, 2005; James et al., 2014; Aldegheri et al., 2023). Therefore, a promising avenue for further alignment of human and machine behavior is to allow machines to interact with their environment (Aubret et al., 2022).

### 5.5. Conclusion

Altogether, our results support previous findings that low shape bias is not a necessary property of convolutional neural networks but is tightly linked to the training data. We show that taking inspiration from human learning strategies over the course of human development can increase shape bias in CNNs whilst simultaneously resolving their trade-off between this bias and performance accuracy. Overall, we show that convolutional architectures which process local spatial information in a brain-like, hierarchical manner align with human classification strategies when training on developmentally inspired images.

## 6. Acknowledgments

This work was funded by the University of Amsterdam Data Science Center Interdisciplinary PhD Program.

## 7. Author contributions

**Niklas Mü ller**: Conceptualization; Data curation; Formal analysis; Investigation; Methodology; Software; Visualization; Writing - original draft; and Writing - review & editing

**Cees G. M. Snoek**: Supervision; and Writing - review & editing

**Iris I. A. Groen**: Conceptualization; Funding acquisition; Methodology; Supervision; and Writing - review & editing

**H. Steven Scholte**: Conceptualization; Funding acquisition; Methodology; Supervision; and Writing - review & editing

## Appendix A. Appendix

### Appendix A.1. Dataset and training information

**Table A.1:** Number of instance in training and test set and number of classes.

|  | ImageNet | MS-COCO | ADE20K | OADS |
| --- | --- | --- | --- | --- |
| Train set | 229111 | 688000 | 507476 | 57496 |
| Test set | 25457 | 86001 | 73129 | 7187 |
| Number of classes | 223 | 80 | 3542 | 16 |

**Table A.2:** OADS object classes.

| OADS Object Class Labels |  |  |  |
| --- | --- | --- | --- |
| Traffic light | *Tree* | *Car* | *Bench* |
| Van | Balcony door | *Scooter* | *Truck* |
| Bollard | Traffic sign | *Bike* | *Bin* |
| SUV | Carrier bike | *Front door* | *Lamppost* |

**Table A.3:**
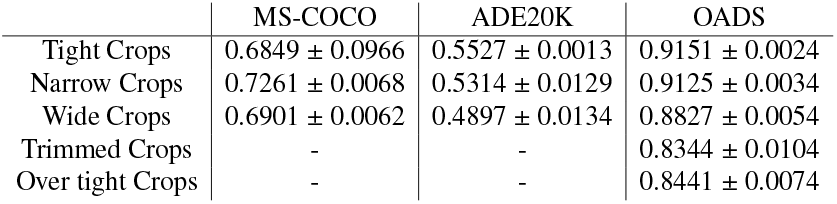
Top-1 test accuracy of ResNet50 after training of 30 epochs.

**Table A.4:**
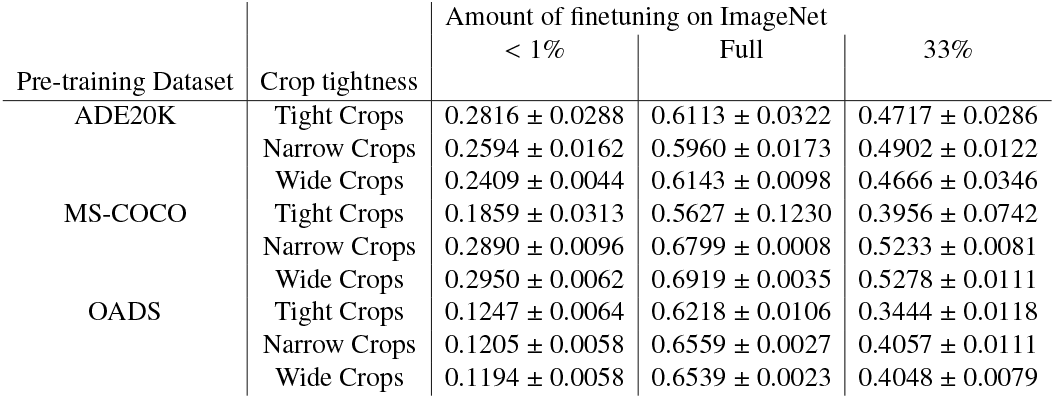
Top-1 test accuracy on ImageNet after finetuning for 15 epochs of ResNet50 pre-trained on various datasets with varying crop tightness.

| Pre-training Dataset | Crop tightness | Amount of finetuning on ImageNet |  |  |
| --- | --- | --- | --- | --- |
|  |  | < 1% | Full | 33% |
| ADE20K | Tight Crops | $0.2816 \pm 0.0288$ | $0.6113 \pm 0.0322$ | $0.4717 \pm 0.0286$ |
| | Narrow Crops | $0.2594 \pm 0.0162$ | $0.5960 \pm 0.0173$ | $0.4902 \pm 0.0122$ |
| | Wide Crops | $0.2409 \pm 0.0044$ | $0.6143 \pm 0.0098$ | $0.4666 \pm 0.0346$ |
| MS-COCO | Tight Crops | $0.1859 \pm 0.0313$ | $0.5627 \pm 0.1230$ | $0.3956 \pm 0.0742$ |
| | Narrow Crops | $0.2890 \pm 0.0096$ | $0.6799 \pm 0.0008$ | $0.5233 \pm 0.0081$ |
| | Wide Crops | $0.2950 \pm 0.0062$ | $0.6919 \pm 0.0035$ | $0.5278 \pm 0.0111$ |
| OADS | Tight Crops | $0.1247 \pm 0.0064$ | $0.6218 \pm 0.0106$ | $0.3444 \pm 0.0118$ |
| | Narrow Crops | $0.1205 \pm 0.0058$ | $0.6559 \pm 0.0027$ | $0.4057 \pm 0.0111$ |
| | Wide Crops | $0.1194 \pm 0.0058$ | $0.6539 \pm 0.0023$ | $0.4048 \pm 0.0079$ |

